# Differential Cross-Protective Immunity is Elicited by Infection with Contemporary Influenza B Lineage Viruses

**DOI:** 10.1101/2024.04.15.589536

**Authors:** Caroline Page, Justin D. Shepard, Sean D. Ray, Jasmine Y. Akinpelu, Ginger Geiger, Stephen M. Tompkins

**Author notes:** Address correspondence to Stephen Mark Tompkins,.

## Abstract

Influenza B virus (FLUBV) significantly contributes to the influenza disease burden and has complicated vaccine development and efficacy, yet remains understudied compared to its counterpart, influenza A virus (FLUAV). Since its isolation in 1940, FLUBV has diverged into two antigenically distinct lineages: Victoria (B/Vic) and Yamagata (B/Yam). Recent human studies and epidemiological modeling reveal differences in immunity elicited by each FLUBV lineage, contributing to higher reinfection rates following B/Yam infection. To investigate disparities in FLUBV lineage cross-protection and immunity, we examined the effects of lineage-specific prior immunity on FLUBV reinfection dynamics. Mice were infected with representative B/Vic and B/Yam viruses from evolutionary distinct clades and subsequently reinfected with heterolineal viruses (i.e., B/Vic → B/Yam and B/Yam → B/Vic) to assess the extent of protection elicited between the lineages. Using this validated challenge model, we explored potential mechanisms underlying the asymmetric reinfection dynamics observed between the lineages. Our findings align with human observations, indicating that contemporary B/Vic viruses confer cross-protection against contemporary B/Yam infections, whereas contemporary B/Yam viruses do not provide the same degree of protection. Furthermore, we demonstrated that serum antibodies elicited by hemagglutinin vaccination cannot account for the observed heterolineal protection. Rather, antibodies targeting the viral neuraminidase (NA) may play a significant role in eliciting cross-protection to subsequent FLUBV infection. Our findings define asymmetric cross-protection resulting from contemporary FLUBV infection and suggest NA as a potential significant contributor to heterolineal FLUBV protection. This asymmetric immunity may also help explain the proposed extinction of B/Yam viruses since the COVID-19 pandemic.

**Importance:** Influenza B viruses (FLUBV) consist of two divergently evolving lineages, Victoria (B/Vic) and Yamagata (B/Yam). Contemporary isolates from these lineages exhibit increased endemic activity and higher evolutionary rates while utilizing distinct mechanisms for evolutionary success. This is exemplified by novel seasonal infection dynamics with Influenza A viruses, differences in cross-protection elicited between the FLUBV lineages, and the potential extinction of B/Yam following the COVID-19 pandemic. We explore FLUBV infection dynamics utilizing contemporary viruses to define the asymmetric immunity elicited between the lineages. Contemporary Yamagata viruses are unable to confer the same breadth of protection as Victoria viruses. This may help explain the higher reinfection rates for Yamagata viruses and suggest a potential contributor to the extinction of this lineage.

## Introduction

The co-circulation of influenza A and B viruses (FLUAV and FLUBV) during seasonal epidemics causes a significant disease and economic burden (1–3). Additionally, it complicates vaccine development and efficacy due to the expansion from bi- and tri-valent formulations to quadrivalent formulations to broaden the breadth of protection from seasonal vaccines (4, 5). Although understudied compared to its FLUAV counterpart, FLUBV plays an essential role in the epidemiology of seasonal infections, accounting for approximately 25% of influenza cases and hospitalizations each season and circulates as the predominant influenza type in one of every seven seasons (3, 4, 6–8). Further, in pediatric patients, the clinical severity of disease is worse for FLUBV infections compared to FLUAV, highlighting the importance of delineating FLUBV infection, reinfection dynamics, and the immune response(s) responsible (3, 6, 7, 9–11). Despite this, significant knowledge gaps persist for FLUBV, and while recent advances have expanded our understanding, considerable progress has yet to be made in unraveling the complexities of FLUBV infection and its impact on host immunity and public health.

Influenza B virus, first identified in the 1940s, subsequently diverged into two antigenically distinct lineages known as Victoria (B/Vic) and Yamagata (B/Yam) in the 1970s (3). Despite regular reassortment events between the two lineages, the separation of the lineages is defined primarily by the hemagglutinin (HA) sequence (12, 13). Both lineages undergo insertions and deletions within the immunodominant surface glycoproteins proteins, HA and neuraminidase (NA), resulting in antigenic drift allowing for viral escape from pre-existing immunity (12). The impact of antigenic drift is remarkable in contemporary post-2015 FLUBV isolates, which have exhibited increased endemic activity, higher evolutionary rates, and multiple genomic selective sweeps, leading to further diversification of each lineage into new immunologically distinct clades and subclades (2, 14, 15). This phenomenon is exemplified with a novel B/Vic subclade, V1A.3, which emerged between 2018-2019, dominated previous subclades, and is connected with an anomalous influenza season in 2019-2020 whereby FLUBV infections circulated early and FLUAV infections rose later in the season (2).

Modeling and retrospective epidemiological studies have begun to investigate group-specific immunity against influenza viruses, with a particular focus on FLUAV and, to a lesser extent, FLUBV (16–18). In humans, homotypic (FLUAV) and intra-lineage (FLUBV) protection from repeat infection occurs with pandemic A/H1N1, A/H3N2, and B/Vic viruses, but not with B/Yam viruses (17). Moreover, a B/Yam infection in one season offers only limited protection against a B/Vic infection in a subsequent season (16). Limited protection from reinfection after exposure to B/Yam viruses observed in humans could potentially justify higher season-to-season reinfections associated with this lineage (4, 16). These observations indicate differences in FLUBV-lineage elicited immunity and suggests the existence of an undescribed phenomenon resulting in reduced protection against reinfection after exposure to B/Yam viruses.

The humoral immune response plays an integral role in influenza cross-protection (19–21). During a naturally acquired influenza infection, antibodies are primarily elicited against the major surface glycoproteins, HA and NA, and are essential in protecting from repeat infections (22, 23). HA-specific antibodies are necessary for viral neutralization and a subset of these antibodies bind moderately conserved epitopes near the globular head and stem, allowing for cross-reactive responses between the FLUAV subtypes and FLUBV lineages (5, 20, 24–26). In addition, NA-specific antibodies have recently been highlighted for playing a role in cross-protective influenza immunity (27–32). While cross-reactive NA-specific antibodies do not prevent infection, they inhibit the enzymatic activity of NA to cleave host cell surface sialic acids to facilitate viral egress, limiting the viral lifecycle, and reducing replication and disease severity (33). Considering the unknown mechanism allowing for persistent reinfection with B/Yam viruses observed in humans, there is a critical need to further investigate the humoral immune response directed towards the HA and NA elicited by FLUBV infection to define their role in this asymmetric immunity.

While human studies have observed intriguing FLUBV infection and reinfection dynamics with contemporary FLUBVs, no experimental animal models reflecting these observations have been established. Here, we established an FLUBV challenge model that explores homolineal (intra-lineage) and heterolineal (inter-lineage) infections in mice to replicate human infection dynamics while utilizing both contemporary (post-2015) and non-contemporary (pre-2015) FLUBV viruses. Our findings reveal that contemporary B/Vic-elicited immunity can provide protection against both homolineal and heterolineal challenges, whereas contemporary B/Yam-elicited immunity is insufficient to fully protect against a heterolineal B/Vic challenge. Utilizing this model, we investigate the mechanisms driving differential immune responses between the FLUBV lineages. We explore the roles of recombinant HA-specific vaccine-elicited immunity and NA-specific infection-elicited immunity in FLUBV cross-lineage protection. These studies further our understanding of the asymmetric FLUBV lineage immunity and allow for a deeper insight to reinfection dynamics.

## Results

### Contemporary FLUBVs elicit asymmetric cross-protection against heterolineal challenge

To assess the immune response to contemporary FLUBV lineages, B/Washington/02/2019 (B/WA) was selected to represent the B/Vic lineage and B/Oklahoma/10/2018 (B/OK) was selected to represent the B/Yam lineage. B/Washington/02/2019 was included in the 2020-2021 quadrivalent influenza vaccine and belongs to the recently emerged subclade, V1A.3, of the Victoria lineage (34). B/Oklahoma/10/2018 (Clade 3A), which is antigenically similar to the B/Yam isolate in the 2020-2021 quadrivalent vaccine, was circulating in the U.S. just prior to the onset of the COVID-19 pandemic. To establish the infectious doses of these viruses in mice, animals were inoculated with a range of viral doses from 10^2^ - 10^5^ PFU and viral replication on day three post-infection was determined via plaque assay of lung homogenates (data not shown). For both B/WA and B/OK, 10^3^ PFU was selected as the infectious dose for our challenge model as this dose caused minimal weight loss and clinical signs yet replicated to similarly high titers in the lungs of mice after infection (Figure 1A and B).

**Figure 1.**
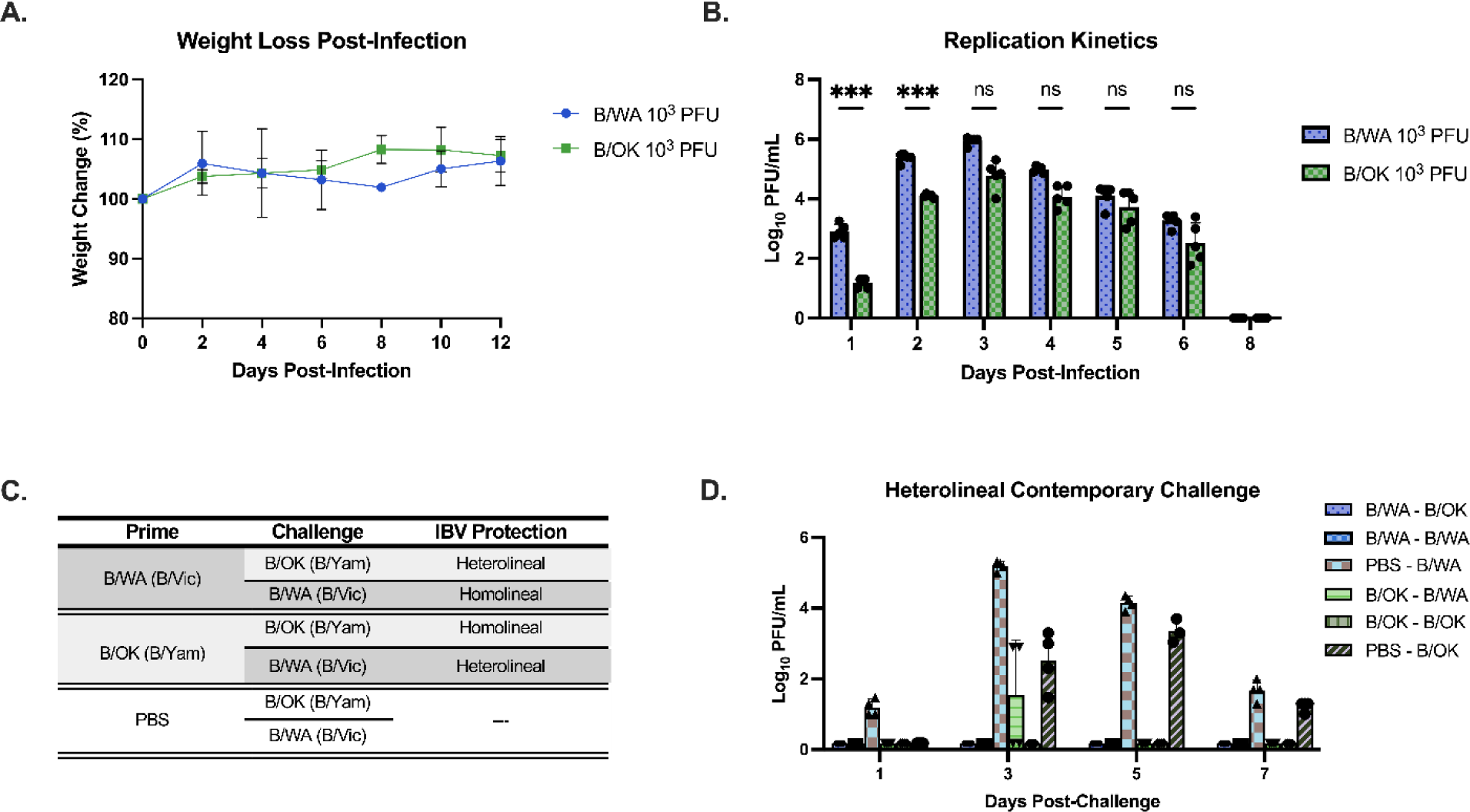
Replication kinetics and breakthrough infections of contemporary FLUBVs. **(A)** Weight loss and **(B)** Lung virus titers of mice infected with 10^3^ PFU of B/Washington/02/2019 (B/WA; Blue) and B/Oklahoma/10/2018 (B/OK; Green). **(C)** Thirty-five days post-infection, mice were challenged with either a homolineal (i.e., B/Yam → B/Yam) or heterolineal (i.e., B/Yam→B/Vic) virus and **(D)** lung virus titers were determined days 1, 3, 5, and 7 post-challenge via plaque assay. Symbols/bars represent mean +/− SD. *P<0.05, Šídák’s multiple comparisons test with n=4 or 5 mice per group

To replicate the human phenomena of limited protection elicited from contemporary B/Yam infection, mice were primarily infected with 10^3^ PFU of B/WA or B/OK. Thirty-five days post-infection (DPI), after clearance of acute infection and establishment of immune memory (35), animals were challenged with a dose of 10^4^ PFU of either a homolineal (i.e. B/Yam → B/Yam) or heterolineal (i.e. B/Yam → B/Vic) virus and lungs were collected on days 1, 3, 5, and 7 post-infection to determine kinetics of viral replication (Figure 1C). All experimental groups were completely protected from reinfection except heterolineal challenged mice infected with B/OK (B/Yam) and subsequently challenged with B/WA (B/Vic, Figure 1D); these breakthrough infections were observed in two-thirds of the animals’ day 3 post-infection across two independent experiments (data not shown). All other homolineal and heterolineal challenged groups were completely protected (Figure 1D). In our mouse challenge model, similar to observations in humans, mice initially exposed to our B/Vic virus displayed complete protection against subsequent B/Yam challenge. However, a majority of the mice that were initially exposed to our B/Yam virus experienced break-through infections on days 3 post-infection, corresponding to the peak of viral replication. There were no clinical signs or weight loss in the experimental groups post-challenge (data not shown). In summary, this mouse challenge model successfully replicates observations in humans that contemporary B/Yam-elicited immunity is unable to afford equal protection from heterolineal reinfection as B/Vic-elicited immunity when utilizing contemporary post-2015 viral isolates.

### Asymmetric cross-protection is a phenomenon not observed with non-contemporary FLUBVs

Considering the notably divergent evolutionary trajectories FLUBV lineages have undergone in recent years (14), we sought to investigate whether the differences in cross-protection between Victoria and Yamagata viruses is restricted to contemporary viruses belonging to recently emerged FLUBV clades or if this phenomenon spans evolutionarily older viruses as well. Representative FLUBV viruses from less contemporary clades were tested using the same timeline as described in Figure 1. Viral replication kinetics for B/Brisbane/60/2008 (B/Bris, Clade 1B) from the B/Vic lineage and B/Massachusetts/02/2012 (B/MA, Clade 2) from the B/Yam lineage were established in mice on days 1, 3, 5 and 7 post-infection with 10^3^ PFU of virus. This infectious dose replicates to similar titers for both viruses each day post-infection and induces minimal clinical signs (Figure 2A and B). To determine whether the evolutionarily older B/Vic-lineage virus can confer cross-protection from a non-contemporary B/Yam heterolineal challenge, mice were primarily inoculated with 10^3^ PFU of either B/Bris or B/MA and challenged thirty-five days later with the heterolineal virus. (i.e. B/Bris → B/MA or B/MA → B/Bris, Figure 2C). In contrast to the contemporary FLUBV challenge (Figure 1), the evolutionarily older B/Vic and B/Yam viruses are able to fully protect from a heterolineal challenge. There was no viral replication in the lungs of heterolineal challenged mice on days 1, 3, 5, or 7 post-challenge (Figure 2D).

**Figure 2.**
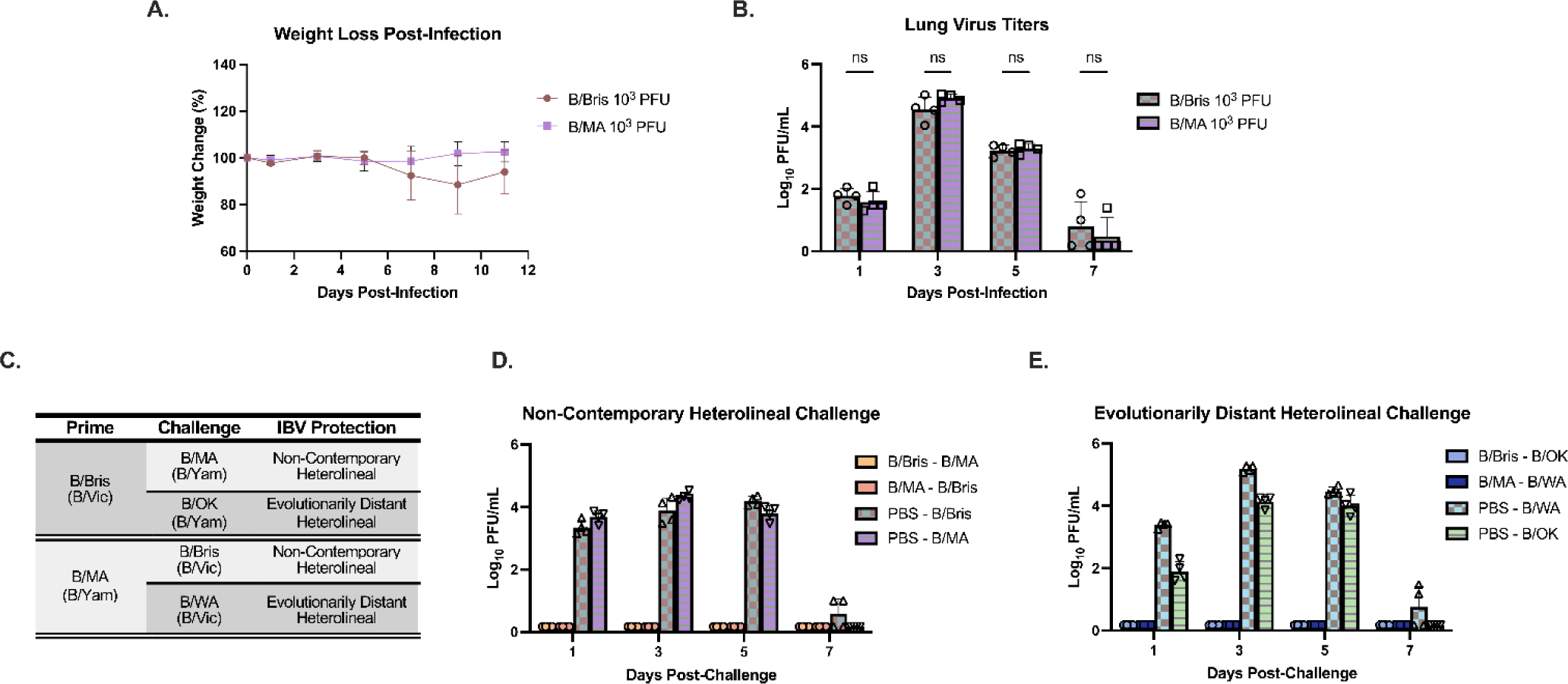
Replication kinetics and breakthrough infections of non-contemporary FLUBV’s. **(A)** Weight loss and **(B)** Lung virus titers of mice infected with 10^3^ PFU of B/Brisbane/60/2008 (B/Bris; brown) or B/Massachusetts/02/2012 (B/MA; purple). **(C)** Thirty-five days post-infection, mice were challenged with a non-contemporary heterolineal virus or an evolutionarily distant heterolineal virus and **(D-E)** lung virus titers were determined on days 1, 3, 5, and 7 post-challenge via plaque assay. Symbols/bars represent mean +/− SD, *P<0.05 one-way ANOVA with n=4 mice per group.

Next, to investigate the ability of parental non-contemporary B/Vic viruses to cross-protect against contemporary B/Yam viruses, and vice versa, animals were inoculated with either B/Bris or B/MA and then challenged thirty-five days later with either B/WA or B/OK (i.e. B/Bris → B/OK or B/Mass → B/WA, Figure 2C). Similar to the non-contemporary challenge with parental strains, evolutionarily older FLUBVs were able to provide cross-protection against the contemporary FLUBVs as demonstrated by the absence of viral replication in experimentally heterolineal challenged mice (Figure 2E). In brief, differential cross-protection observed with the contemporary FLUBV isolates is a novel phenomenon that did not occur with prior FLUBV clades.

### Passive transfer of FLUBV anti-sera confers differential cross-protection

Next, we assessed whether passive transfer of anti-sera elicited from a contemporary FLUBV infection would be sufficient to replicate protection dynamics from homo- or heterolineal challenge as observed in Figure 1. Mice were infected with 10^3^ PFU of either B/WA (B/Vic) or B/OK (B/Yam) to elicit virus-specific serum antibodies. Thirty-five days post-infection, serum was collected and transferred to naïve recipient mice who were then challenged with either a homolineal or heterolineal virus (B/WA or B/OK). Victoria-elicited antibodies significantly reduced viral replication following homolineal B/Vic challenge (Figure 3A) as well as heterolineal B/Yam challenge (Figure 3B). Conversely, while Yamagata-elicited antibodies significantly reduced viral replication against a homolineal B/Yam challenge (Figure 3B), they failed to protect against heterolineal B/Vic challenge (Figure 3A). Contemporary Victoria lineage specific antibodies elicited from infection can provide protection from both homo- and heterolineal challenges while B/Yam-elicited antibodies are insufficient in providing heterolineal protection. The asymmetrical reinfection dynamics observed in Figure 1 were recapitulated in this passive transfer experiment, demonstrating serum antibodies are mediating FLUBV heterolineal immunity.

**Figure 3.**
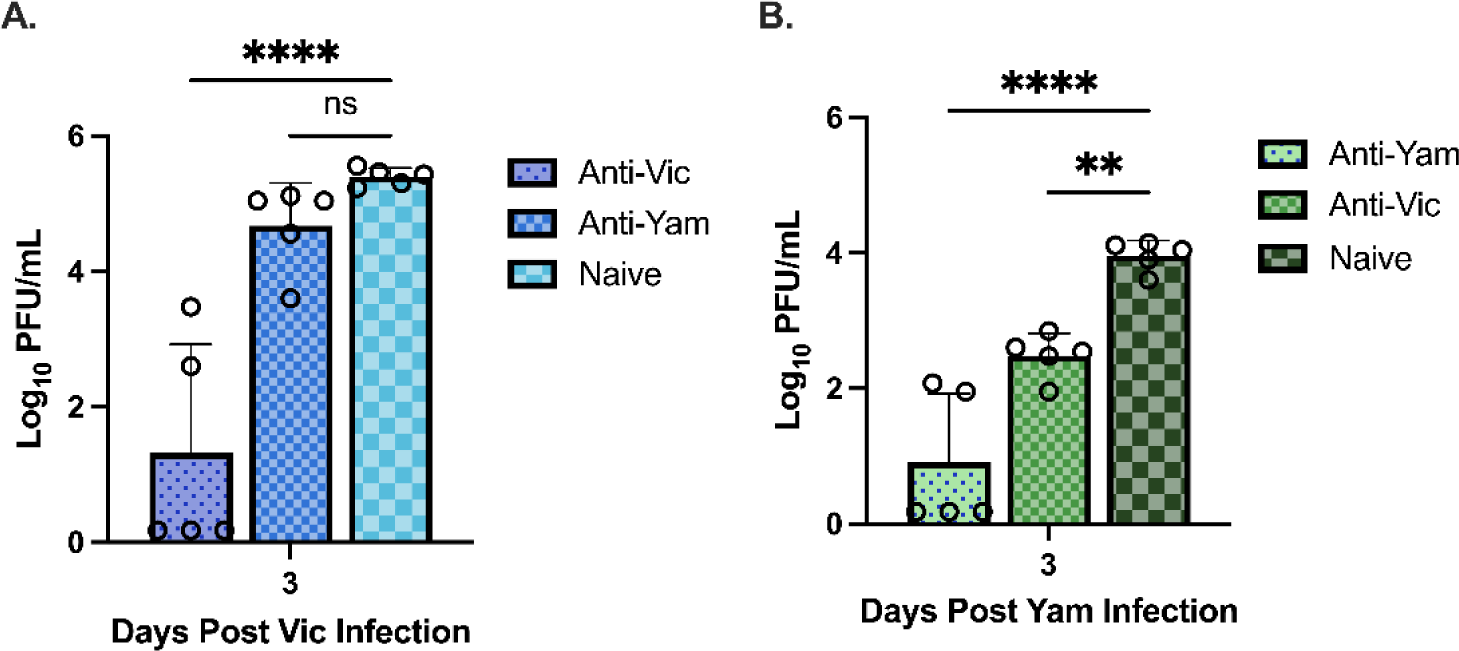
Passive transfer of polyclonal FLUBV antibodies confer cross-protection. Lung virus titers in mice receiving Anti-Victoria (Anti-Vic; B/Washington/02/2019) antibodies or Anti-Yamagata (Anti-Yam; B/Oklahoma/10/2018) antibodies three days following an infection with **(A)** B/Washington/02/2019 (Victoria; blue) or **(B)** B/Oklahoma/10/2018 (Yamagata; green). Data is representative of n=5 mice per group. Symbols/bar represent the mean +/− SD. *P<0.05, one-way ANOVA.

### HA-specific antibodies are insufficient to confer cross-lineage protection

The defined divergence between the FLUBV lineages is based on sequence homology of the HA protein. Therefore, observations of differential immunity in our mouse model led to us hypothesize that HA-specific antibodies alone are sufficient in driving differential cross-protection. To test this, we immunized mice with either recombinant HA of B/Washington/02/2019 (Vic-rHA) or B/Oklahoma/10/2018 (Yam-rHA) in a prime only, or homolineal/heterolineal prime-boost regimen (Figure 4A). Two weeks post-boost, serum was collected, and animals were challenged with 10^4^ PFU of either B/WA or B/OK. Cross-reactive IgG antibodies were elicited following a single rHA immunization and titers were boosted following the second immunization (Figure 4B and C). Hemagglutination inhibition titers showed no cross-reactivity and only the B/Yam Prime – B/Yam Boost group displayed titers greater than 40 (data not shown). Following challenge, the homolineal prime-boost regimens (i.e. Vic-rHA → Vic-rHA or Yam-rHA → Yam-rHA) significantly reduced viral replication against the homolineal challenges for both B/Vic and B/Yam lineages (Figure 4D and E). Additionally, the rHA B/Vic prime – B/Yam boost significantly reduced viral replication from a B/Yam challenge (Figure 4E), and while the same trend was observed for the rHA B/Yam prime – B/Vic Boost against the B/Vic challenge, it was not significant (Figure 4D). Although homolineal protection is conferred by homolineal rHA prime-boost induced HA-specific antibodies, heterolineal cross-protection was not induced from rHA immunization alone. In our mouse infection model (Figure 1), we observed cross-lineage protection afforded by B/Vic, but not B/Yam infection; here, both B/Vic-rHA and B/Yam-rHA immunizations were insufficient in providing cross-lineage protection, suggesting differential immunity observed following FLUBV infection cannot be explained by HA-specific antibody responses.

**Figure 4.**
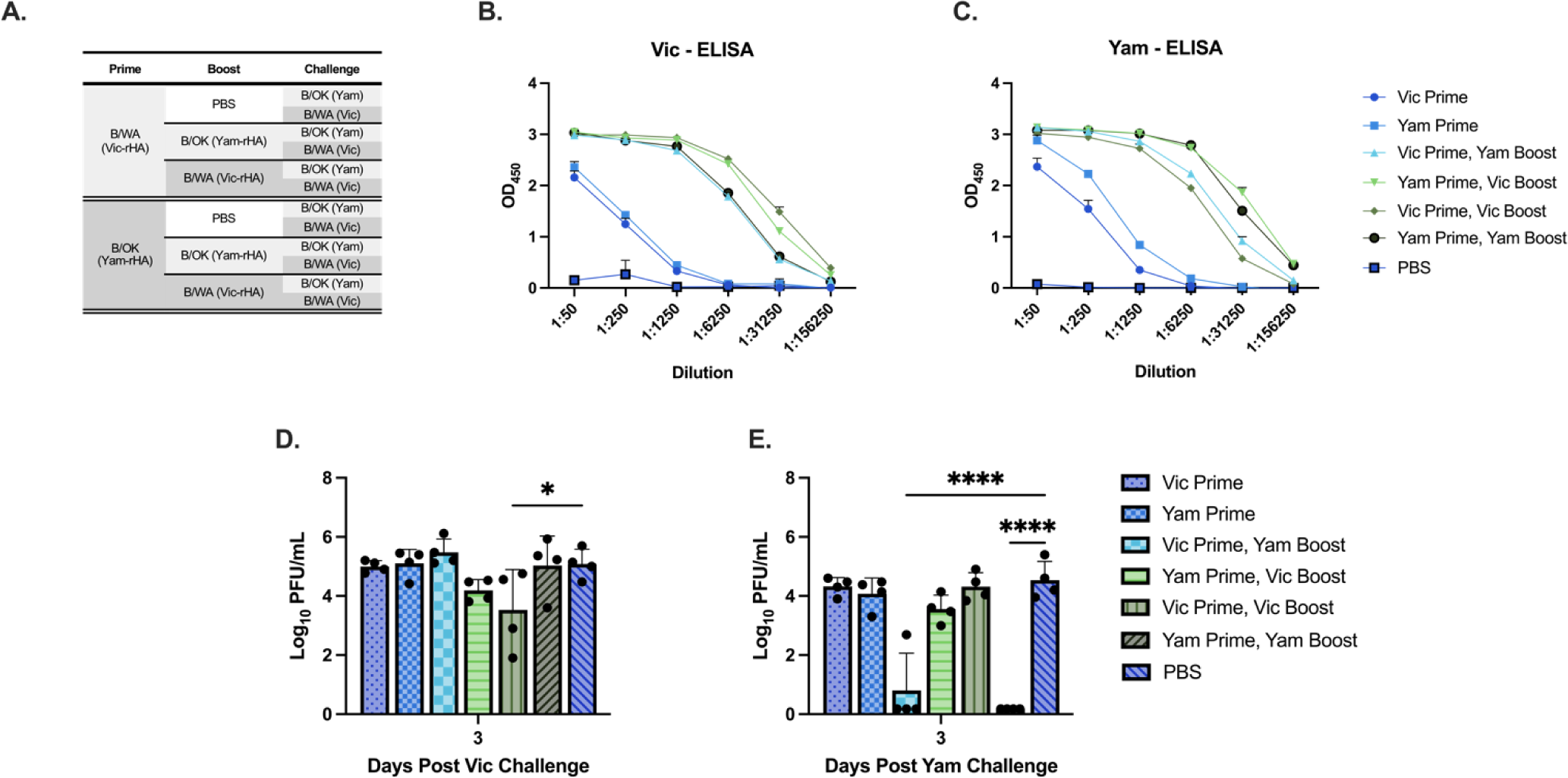
HA-specific antibodies are insufficient to confer cross-lineage protection. (**A**) Vaccination groups and challenge viruses **(A-B)** IgG serum antibody titers measured by ELISA fourteen days post-boost against **(B)** B/Victoria recombinant HA protein (B/Washington/02/2019) and **(C)** B/Yamagata recombinant HA protein (B/Oklahoma/10/2018). Data is representative of the average OD value from n=4 mice per group. **(D-E)** Lung virus titers three days after FLUBV challenge with **(D)** B/Washington/02/2019 (Vic) and **(E)** B/Oklahoma/10/2018 (Yam). Symbols/bars represent mean +/− SD of n=4. *P<0.05, one-way ANOVA.

### Cross-reactivity of FLUBV NA-specific immune serum contributes to asymmetric immunity

To elucidate the role of cross-reactive antibody responses elicited by heterolineal FLUBV infection, we examined systemic HA- and NA-specific serum antibody responses in mice thirty-five days post-FLUBV infection. Hemagglutination inhibition assays (HAI), considered the gold standard for assessing protection against influenza infection (36), were employed to evaluate the HA-specific responses. Our findings revealed serum HAI titers against the homologous virus were higher than those against the heterolineal antigen, exhibiting minimal cross-reactivity (Figure 5A). Notably, the non-contemporary viruses, B/MA and B/Bris, conferred broader overall HAI titers compared to the contemporary viruses. Influenza B/MA (B/Yam) elicited anti-serum exhibited cross-reactive HAI titers to the homolineal B/OK (B/Yam), whereas B/OK-serum had antibody titers exclusively towards the homologous (B/OK) antigen. Meanwhile, B/Bris (B/Vic) elicited serum was the sole antiserum demonstrating cross-reactive HAI titers, albeit weak, towards the heterolineal B/MA (B/Yam) antigen.

**Figure 5.**
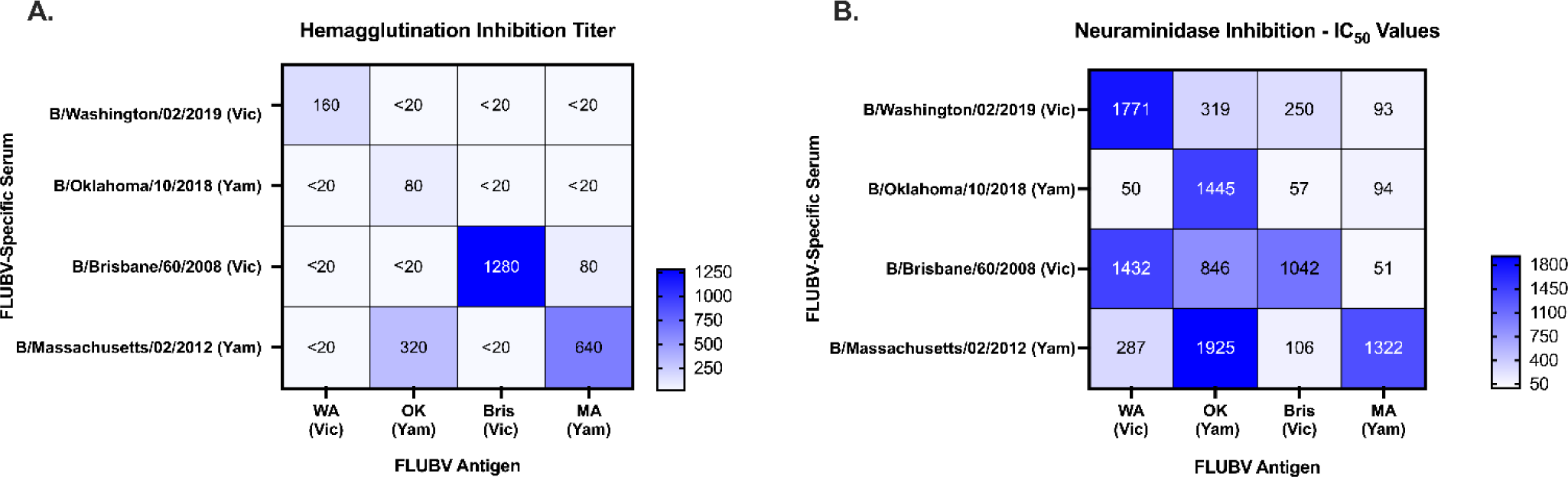
Cross-reactivity of FLUBV-immune serum. Thirty-five days post-infection with B/Washington/02/2019, B/Oklahoma/10/2018, B/Brisbane/60/2008, or B/Massachusetts/02/2012 serum was collected and assessed via hemagglutination inhibition assays (HAI) and enzyme-linked immunosorbent assay (ELLA). **(A)** FLUBV-specific serum (y-axis) were tested against homolineal or heterolineal antigens (x-axis) to determine cross-reactivity of the HA response. The HI titers, averages from n=4 mice, are reported as the reciprocal dilution that resulted in inhibition of agglutination. **(B)** Neuraminidase inhibition as determined by ELLA assay. The values reported are averages of n=4 animals IC_50_ values determined from linear regressions.

In addition to evaluating HAI antibody responses, we also assessed NA-inhibiting (NAI) antibody responses as NAI antibodies have been associated with protection from infection (28, 37). All FLUBV infection elicited robust NAI antibody responses to the homolineal antigens (Figure 5B). Notably, the contemporary B/Yam virus (B/OK) induced the least cross-reactive NA-specific antibodies compared to the other FLUBVs. Conversely, the contemporary B/Vic virus (B/WA) elicited cross-reactive NA-specific antibodies to both the homolineal B/Bris and heterolineal B/OK viruses. The non-contemporary B/Bris and B/MA viruses exhibited superior efficacy compared to our contemporary viruses in eliciting homo- and heterolineal cross-reactive NA-specific antibody responses.

In summary, FLUBV infections with contemporary and non-contemporary viruses resulted in limited cross-reactive HAI titers. Conversely, NA-specific antibody responses exhibited a degree of both homo- and heterolineal cross-reactivity by ELLA, with the exception of the contemporary B/Yam virus, B/OK, suggesting a significant role for NA in contemporary FLUBV infection-elicited asymmetric immunity.

## Discussion

Influenza B virus infection is a significant contributor to the overall influenza disease and economic burden, but remains understudied compared to FLUAV (6). Reinfection dynamics between the lineages have been previously modeled and cross-reactive humoral responses from both natural infection and seasonal vaccination have been described (21, 27, 38). However, to our knowledge, no studies have assessed homolineal (intra-lineage) or heterolineal (inter-lineage) protection following natural infection in animals utilizing both contemporary and non-contemporary viruses. In this study, using a mouse model, we recapitulated human observations of contemporary FLUBV reinfection dynamics, which suggest B/Yam-elicited immunity is insufficient in conferring protection from a cross-lineage B/Vic (heterolineal) infection (16, 17, 38). We show that animals primarily infected with contemporary B/Vic virus are protected from a contemporary heterolineal (i.e. B/Vic → B/Yam) challenge. In contrast, animals primarily infected with contemporary B/Yam virus were susceptible to reinfection when exposed to a contemporary heterolineal (i.e. B/Yam → B/Vic) challenge. These data suggest there exists differential infection dynamics and subsequent elicited immunity between contemporary FLUBV lineages which allows for B/Vic-elicited cross-lineage protection against B/Yam, but incomplete B/Yam-elicited cross-protection against B/Vic. In addition, utilizing less contemporary FLUBV isolates, we show that this previously undescribed phenomena of asymmetric protection are observed only with the contemporary post-2015 FLUBV viruses; whereas non-contemporary B/Vic and B/Yam viruses fully protected mice from heterolineal FLUBV reinfection independent of the challenge virus clade.

We established an animal model to define cross-protection elicited from FLUBV infection and investigated potential immune mechanisms responsible for the differential responses observed between the contemporary FLUBV lineage viruses. We first show the mechanism of protection is antibody-mediated with the passive transfer of infection-elicited immune serum. When antibodies obtained from contemporary FLUBV infection were passively transferred into naïve recipients and challenged with the homo- or heterolineal viruses, we found that B/Yam antibodies alone provide less protection against a B/Vic challenge compared to B/Vic antibodies against B/Yam (Figure 3). Based on previous studies, we initially hypothesized the differences observed in cross-protection could be explained by cross-reactive humoral responses directed at the globular head or stem domain of the hemagglutinin (21, 39). However, vaccination with recombinant HA failed to elicited cross-protection from a heterolineal challenge (Figure 4). Homologous B/Vic and B/Yam prime-boost groups were only capable of eliciting protection against a homolineal challenge, indicating that HA-specific immunity, neutralizing or non-neutralizing, was insufficient to explain the differences in cross-protection observed with contemporary FLUBV infections.

In our studies, we deliberately chose infectious doses that induced minimal to no clinical signs. However, the noticeable decrease in viral replication and the more rapid viral clearance observed in animals with break-through infections when compared to the control animals prompted us to expand our studies to evaluate the influence of the NA-specific antibody response on cross-protection against FLUBV. Although NA antibodies have been implicated in broad protection against heterologous strains of FLUAV, the role for infection-elicited cross-lineage protective NA antibodies has not been fully defined for FLUBV (22, 30). Neuraminidase-specific antibodies function by binding to the NA protein and blocking its enzymatic activity preventing the virus from being released by infected cells and limits the ability of the virus to move within the respiratory tract (33). Hence, when NA-specific antibodies are involved in protection from infection, it can clinically result in reduced disease severity and duration of viral shedding, rather than complete viral neutralization which is associated with HA-mediated protection (29, 37). Unlike frequent HA mutations, the NA protein for FLUBV is under less immune pressure, has fewer amino acid mutations, and remains antigenically analogous between FLUBV strains (14, 33, 40). In contrast to the HA-specific response, which was mostly restricted to homologous antigens, the NA-specific antibodies elicited from FLUBV infection demonstrated cross-reactivity across lineage and evolutionary clade except for the contemporary B/Yam virus which was more restricted in the NA-specific response (Figure 5B). We suggest this restricted NA response contributes to the insufficient protection elicited from contemporary B/Yam infections.

Interestingly, the unique reinfection dynamics of B/Yam, resulting in breakthrough infections, were observed exclusively with our contemporary FLUBV viruses. The viral isolates used in these studies were selected based on the evolutionary clade to which they belong (14). The less contemporary viruses, B/Bris and B/MA, belong to older clades, while B/WA and B/OK belong to newer clades. Since 2015, contemporary viral isolates from each lineage exhibited higher levels of endemic activity and have employed distinct mechanisms for evolutionary success (14). The B/Vic viruses evolved through major changes to the HA protein, while B/Yam viruses experienced stronger selective pressure towards the NA protein (14). The distinct evolutionary pressures between the lineages in recent years, coupled with limited reassortment events, have led to a further diversification between the lineages. This contrasts with less contemporary viruses, where historically frequent reassortment has been observed between the lineages. For example, in the early 2000s, the NA gene segment from the Victoria lineage viruses was replaced by the Yamagata lineage NA, introducing conserved epitopes in the B/Vic lineage that could elicit cross-reactive antibody responses to B/Yam lineage viruses (41). This could explain the cross-protection that occurred with the less contemporary viruses that was not observed with the contemporary viruses.

While a comprehensive assessment of the immune response to contemporary FLUBV viruses is incomplete, we suggest that the observed lineage differences in this study between contemporary and their parental, non-contemporary viruses are, in part, due to the recent further divergence of the lineages. This divergence results in less cross-reactivity between the contemporary viruses as they are more antigenically distinct. However, our contemporary B/Vic infection maintained the ability to elicit NA-specific antibodies that interfered with both B/Vic and B/Yam infection, while our contemporary B/Yam infection elicits NA-specific antibodies that only prevent B/Yam infection. We suggest the differences between the contemporary lineage infections occurs in part because B/Yam lineage viruses experienced significant NA antigenic drift (14), resulting in a restricted antigen specificity for contemporary B/Yam FLUBV-specific immunity, allowing for breakthrough infections and greater endemic activity for B/Vic viruses. Additionally, in recent years, B/Yam viruses have displayed increased relative genetic diversity of the HA and NA proteins, with peaks in 2015 and 2018 (14). Our B/Yam virus, B/Oklahoma/10/2018, is representative of this new clade 3A genetic diversity which is characterized by NA substitutions at I171M and N342K. While previous studies show differences in NA enzymatic activity and FLUBV pathogenicity based on substitutions in the NA at position 342, the impact this has on subsequent heterolineal cross-protection is unknown(42, 43). We suggest these B/Yam contemporary mutations could explain why the NA-specific antibody responses were uniquely restricted to the homologous antigen. This restricted immune profile could, in turn, allows for the B/Vic breakthrough infections to occur, as limited cross-reactive antibodies are elicited by B/Yam infection. If B/Vic infections continued to induce cross-protection to both lineages, while B/Yam infections only conferred protection against B/Yam, there would exist a population with greater immunity against B/Yam viruses compared to B/Vic viruses. These dynamics could help explain the disappearance of the Yamagata lineage following the COVID-19 pandemic as B/Yam viruses had a reduced population of susceptible individuals (4, 44).

## Methods

### Virus Propagation

The FLUBVs B/Washington/02/2019 (International Reagent Resource (IRR), FR-1709, E6), B/Oklahoma/10/2018 (IRR, FR-1660, C2/E1), B/Brisbane/60/2008 (St. Jude Children’s Research Hospital, E4), and B/Massachusetts/02/2012 (IRR, FR-1196, E6) were propagated in 9-to-11-day-old specific pathogen free embryonated chicken eggs. Allantoic fluid was inoculated, and eggs were incubated at 35°C with 5% CO_2_ for 3 days. After embryo termination in 4°C overnight, allantoic fluid was collected and clarified by centrifugation at 4000 x *g* for 20 minutes and stored at −80°C until further use.

### Mice, Infections, and Immunizations

Six- to eight-week-old, female BALB/c mice were purchased from Charles River and utilized for all experiments. For infections, mice were anesthetized using 5% isoflurane and intranasally inoculated with 30μL of live virus at specified infectious doses. Mice were monitored daily for weight loss and clinical signs. Animals reaching institutionally defined humane endpoints were humanely euthanized in accordance with American Veterinary Association Guidelines. For immunizations, each mouse was intramuscularly injected with 50μL of a 2μg dose of recombinant protein diluted 1:1 with AddaVax (InvivoGen, Cat # vac-adx-10) in the hind leg. Experiments were performed in accordance with protocols approved by the University of Georgia Institutional Animal Care Committee (IACUC).

### Sample Collection

Lung samples were collected into 1mL of phosphate-buffered saline (PBS) at specified timepoints, mechanically homogenized, and clarified by centrifugation at 2500 x *g* for 2 minutes. Supernatant was collected, aliquoted, and stored at −80°C until quantification via plaque assay. Blood was collected in blood collection tubes (BD Microcontainer, Ref #365967) at specified timepoints by submandibular or terminal bleeds and centrifuged at 10,000 x *g* for 2 minutes to separate serum. Serum was aliquoted and stored at 4°C until further use.

### Homo- and Heterolineal Challenge Model

To establish infectious doses, groups of 4 mice were intranasally infected with B/Washington/02/2019 (B/WA), B/Brisbane/60/2008 (B/Bris), B/Oklahoma/10/2018 (B/OK), or B/Massachusetts/02/2012 (B/MA) at doses ranging from 10^2^ – 10^5^ PFU. On day 3 post-infection, animals were sacrificed, and lungs were collected for assessment of viral replication via plaque assay. Weights were monitored daily until full recovery. For replication kinetics experiments, animals (n=4 per infection) were intranasally infected with either 10^3^ PFU of B/WA, B/Bris, B/OK or B/MA. Lungs were collected at specific timepoints post-infection for quantification of viral replication.

For the homo- and heterolineal challenge infections, groups of 4 mice were primarily inoculated with an infectious dose of 10^3^ PFU of either the representative B/Victoria virus (B/Washington/02/2019 or B/Brisbane/60/2008) or B/Yamagata virus (B/Oklahoma/10/2018 or B/Massachusetts/02/2012) and monitored daily for clinical signs. Thirty-five days post-infection, animals were challenged with a dose of 10^4^ PFU of either the homolineal or heterolineal virus. Lungs were collected on day 1, 3, 5, and 7 post-challenge (DPC) for assessment of viral replication. Weights and clinical signs were monitored daily post-challenge.

### Recombinant Protein Vaccination

Groups of 4 mice were primarily vaccinated with recombinant protein or PBS on day 0 and boosted with PBS, homologous, or heterologous FLUBV recombinant protein 21 days post-vaccination (DPV 21). On DPV 21 and 35, serum was collected for serological analysis. Animals were challenged with 10^4^ PFU of B/WA or B/OK and lungs were collected DPC 3 for viral quantification.

### Serum Passive-Transfer

Groups of 15 mice were inoculated with PBS or 10^3^ PFU of B/WA or B/OK and terminally bled thirty-five days post-infection for serum collection and serological analysis. Serum was inactivated in a 56°C heat bath for 45 minutes. Individual animals’ serum was then assessed for antigen specificity using whole-virus ELISAs. Once reactivity was confirmed, serum from each group was pooled and 250μL of inactivated serum was transferred by intraperitoneal (I.P) injection into naïve recipient mice (n=10 per group). Immediately following transfer, groups were challenged with 10^3^ PFU of either B/WA or B/OK. Lungs were collected three days post-challenge for assessment of viral replication.

### Plaque Assay

Madin-Darby Canine Kidney (MDCK) cells were cultured in Dulbecco’s Modified Eagle Medium (DMEM, Corning, Ref #10-013-CV) supplemented with 5% fetal bovine serum, 1x antibiotics-antimycotic (GenDEPOT, Cat # 002-010) and incubated at 37°C with 5% CO_2._ To determine virus titers, MDCK cells were seeded with 4×10^5^ cells per well in a 12-well tissue culture treated plate (Thermo Scientific, Ref #130185) and incubated for 24 hours at 37°C with 5% CO_2._ Cells were then washed twice with 1x sterile PBS (Corning). Tenfold serial dilutions of virus stocks were prepared in DMEM with 0% FBS. MDCK cells were then inoculated with dilutions and incubated for 1 hour at 35°C with 5% CO_2._ Following incubation, the inoculum was overlayed with 0.8% avicel and MEM supplemented with 1M HEPES, 200mM L-glutamine, 7.5% NaHCO3 and 1X antibiotics-antimycotics and incubated at 35°C with 5% CO_2_ for 72 hours. Following incubation, overlay was decanted, and wells were washed twice with 1x PBS, then fixed with methanol:acetone (80:20). Subsequently, wells were stained with crystal violet, and plaques were counted to determine titers in plaque forming units per mL (PFU/mL).

### Enzyme-Linked Immunosorbent Assay (ELISA)

Ninety-six well microtiter plates (Greiner bio-one, Ref #655001) were coated with 100μL of a 1:100 dilution of either live B/WA or B/OK or recombinant protein at 1ug/mL and incubated overnight at 4°C. Plates were then washed 3x with 0.05% PBS-T (PBS and 0.05% Tween-20) and blocked with 200μL of 3% non-fat dry milk for 2-hours at room temperature (RT). Serum was inactivated at 56°C for 45 minutes and serially diluted ten-fold in blocking buffer then incubated for 2-hours at RT. Following incubation, plates were washed 3x and HRP-conjugated goat anti-mouse IgG secondary antibody (Thermo, Cat #31430) was diluted 1:3500 in blocking buffer and added to all wells (100μL/well). Following a 1-hour incubation at RT, plates were washed 3x and 50μL of TMB-substrate solution (Vector Laboratories, Inc. SKU-4400) was added to all wells for 10 minutes. The reaction was stopped with 50μL of 1N H_2_SO_4_ and was read at a wavelength of 490 nm using Cytation 7 (BioTek).

### Hemagglutination Inhibition Assay (HAI)

To remove non-specific inhibitors of HA, serum was treated with a 1:4 dilution of receptor-destroying enzyme (RDE) and incubated overnight (16-18 hours) at 37°C followed by a 30-minute inactivation at 56°C and subsequent dilution to 1:10 with PBS. Prior to starting the assay, viral stocks were diluted to 8 hemagglutinating units (HAU) per 50 μL (4 HAU/25μL). RDE-treated serum was added to a 96-well V-bottom plate, serially diluted 2-fold (starting from 1:2) and incubated for 30 minutes at RT after the addition of 25μL of 4 HAU of virus.

Subsequently, 50μL of 0.5% turkey red-blood cells were added and the plate was incubated for 30 minutes at RT. Following the final incubation, the plate was tilted, and results were recorded as the reciprocal of the highest dilution that prevented the virus from agglutinating.

### Enzyme-Linked Lectin Assay (ELLA)

To determine viral NA activity, maxisorp surface 96-well microtiter plates were coated with 25 µg/mL of fetuin (Sigma CAT#F3004-25MG) in coating buffer (PBS) and incubated at 4°C overnight. The following day, plates were washed 3x with PBS-T wash buffer (0.05% Tween 20). Virus was serially diluted 2-fold and 50μL of viral dilutions along with 50μL of sample diluent (PBS-Ca/Mg, 1% BSA, 0.5% Tween 20) were added to the plate and incubated at 37°C with 5% CO_2_ for 18 hours. After incubation, plates were washed 6x and 100μL of PNA-HRP (1:1000 dilution) was added to all wells for a 2-hour incubation at room temperature. Plates were washed 3x and developed with TMB-substrate solution (Vector Laboratories, Inc. SKU-4400) for 15 minutes. The reaction was stopped with 100μL of 1N sulfuric acid, and the optical density was read at 450nm using Cytation 7 (Biotek).

To perform NI assays, plates were coated as described above. Heat inactivated serum was diluted 2-fold in PBS at a starting dilution of 1:10. 50μL of previously determined optimized virus dilution was added to 50μL of serum dilutions. Fetuin coated plates were washed 3x with PBS and 100μL of the virus:serum mixture was added to plates which were incubated at 37°C for 18 hours. After incubation, plates were washed, developed, and read as described above.

### Plasmids, Protein Expression and Purification

The recombinant B/Washington/02/2019 HA was kindly provided by Dr. Ted Ross (University of Georgia). The ectodomain of influenza B/Oklahoma/10/2018 HA was cloned into a pCAGGs plasmid vector. Plasmids were transformed into E. coli (DH5a, NEB), grown to an OD_600_ of 0.6 in Luria Broth, and extracted using a Zymo II Maxi Prep kit according to the manufacturers protocol (Zymogen). For transfections, suspension cultures of expi293F cells (ThermoFisher) grown at 37°C and 8% CO_2_ were prepared and transfected with plasmids according to the manufacturers protocol (ThermoFisher). Briefly, 1µg of plasmid DNA per mL of expi293F cells were complexed with Expifectamine transfection reagent (ThermoFisher) and incubated at room temperature for 20 minutes. Expi293F cells with >95% viability were diluted to 3×10^6^ cells/mL, and the DNA was added to the culture flask, and returned to incubation. Enhancers were added according to the manufacturers protocol 18 hours post-transfection (ThermoFisher). Following 5-7 days of incubation at 37°C and 8% CO_2_, cells were pelleted by centrifugation at 3,000 x *g*, and supernatants were vacuum filtered using a 0.2 μm membrane filter. The filtered supernatant was then applied to a nickel agarose resin (ThermoFisher) packed gravity flow column. Five column volumes of wash buffer [20mM Tris HCl, 500mM NaCl, 20mM imidazole, pH 7.5] was applied to the resin before eluting the protein with 5 column volumes of elution buffer [20mM Tris HCl, 500mM NaCl, 250mM Imidazole]. The protein was then concentrated in a 30K MWCO centrifugal filter, centrifuged at 3,000 x g (ThermoFisher).

## Acknowledgments and funding sources

This project has been funded in whole or in part with Federal funds from the National Institute of Allergy and Infectious Diseases, National Institutes of Health, Department of Health and Human Services, under Contract No. 75N93021C00018 (NIAID Centers of Excellence for Influenza Research and Response, CEIRR). Madin-Darby Canine Kidney (MDCK-ATL) Cells, FR-926, as well as FLUBVs B/Washington/02/2019 (Victoria Lineage; FR-1709), B/Massachusetts/2/2012 (Yamagata Lineage; FR-1196), and B/Oklahoma/10/2018 (NA D197N) (Yamagata Lineage; FR-1660) were obtained through the International Reagent Resource, Influenza Division, World Health Organization Collaborating Center for Surveillance, Epidemiology and Control of Influenza, Centers for Disease Control and Prevention, Atlanta, Georgia, USA.

## Author Contributions

C.P. and S.M.T designed research; C.P., J.D.S., S.D.R., G.G. and J.A. performed research; C.P., and S.M.T. analyzed data; and C.P. and S.M.T. wrote the paper.

